# Combination of mitochondrial tRNA and OXPHOS mutation reduces lifespan and physical condition in aged mice

**DOI:** 10.1101/233593

**Authors:** G. Reichart, J. Mayer, T. Tokay, F. Lange, C. Johne, S. Baltrusch, M. Tiedge, G. Fuellen, S. Ibrahim, R. Köhling

## Abstract

Mutations in the mitochondrial DNA (mtDNA) are widely known to impact on lifespan and tissue integrity. For example, more than 250 pathogenic mtDNA mutations are known, many of which lead to neurological symptoms. In addition, major neurodegenerative diseases share key components of their etiopathogenesis with regard to mtDNA mutations, mitochondrial dysfunction and oxidative stress. In our study we used a set of conplastic mouse models carrying stable point mutations in mitochondrial genes of transfer RNA (tRNA) and oxidative phosphorylation (OXPHOS)-proteins. We analyzed the impact of these mutations on complex traits like lifespan, learning and memory in the ageing process. The combination of both point mutations in the OXPHOS complex IV gene and adenine insertions in the mitochondrially encoded tRNA arginine (tRNA-Arg) gene (*mt-Tr*) leads to an age-dependent phenotype with elevated mitochondrial superoxide production in the neocortex. Mice with this combination of tRNA and OXPHOS mutations show significantly reduced lifespan and poor physical constitution at the age of 24 months, whereas single point mutations in OXPHOS or mt-tRNA(Arg) do not have this impact. Therefore, we suggest a synergistic effect of these mutations.

## Introduction

Mitochondria represent more than just the major energy production site of our cells. They play a crucial role in cell homeostasis [1], apoptosis [2], cellular stress responses [3], nuclear gene expression [4], immune response [5] and even synaptic transmission [6]. Their genome (mitochondrial DNA; mtDNA) encodes for critical mitochondrial components, i.e. genes coding for oxidative phosphorylation (OXPHOS) protein subunits, tRNAs and rRNAs [7]. Mutations in mtDNA reportedly cause mitochondrial dysfunction and disease phenotypes. Already decades ago, first reports showed that point mutations / deletions in mtDNA can provoke severe diseases, e.g. Leber’s hereditary optic neuritis and mitochondrial myopathy, in humans [8, 9]. Since then, heritable pathologies have no longer been exclusively attributed to the nuclear genome. MtDNA variants / mutations have also been associated, for example, with susceptibility to common diseases like type 2 diabetes [10]. As the brain, being to a large extent post-mitotic, is particularly vulnerable to mtDNA mutations [11] and mitochondrial dysfunction may be due to a higher possibility of clonal expansion of mutated mtDNA [12], it is not surprising that mtDNA dysfunction can be identified in some of the most prevalent neurological disorders among adults [13, 14]. The pathogenesis of Parkinson’s disease (PD), for instance, is associated with mtDNA deletions [15] and mitochondrial dysfunction [16]. PD, Alzheimer’s disease (AD) [17] and amyotrophic lateral sclerosis (ALS) [18] are major neurodegenerative diseases which share key components of their etiopathogenesis in view of either mtDNA mutation, mitochondrial dysfunction or oxidative stress [19]. Furthermore, a compromised shape of mitochondria has been described as a biomarker for neurological disease in general [20], and it is thought to be linked to detrimental mitochondrial dynamics - i.e. fission and fusion [21, 22].

The diversity and high frequency of mtDNA mutations highlights the necessity of investigating the genetics of mitochondrial dysfunction in depth [23]. The progression of mitochondrial disease is known to vary, depending on the level of heteroplasmy, causing more severe characteristics under outbalance of mutant mtDNA [24, 25]. Possibly, there are also other mechanisms influencing the gap between genotype and phenotype, among which heteroplasmic transcriptional reprogramming was proposed [26].

Remarkably, most of the mtDNA point mutations described apply to mitochondrial tRNA mutations [27]. Point mutations in mitochondrial tRNA genes have been shown to modulate mitochondrial function, and have been linked to several syndromes and grave pathological conditions (e.g. myopathy, encephalomyopathy, ataxia and multi-organ failure) [28]. Specifically, a point mutation in the gene coding for mt-tRNA(Asp) is associated with multisystemic mitochondrial disease [29]. Further, mutations in mt-tRNA(Arg) were reported to result in mitochondrial encephalomyopathy [30, 31]. Lastly, a mt-tRNA(Lys) mutation was related to mitochondrial dysfunction associated with myopathy and exercise intolerance [32].

However, these insights from human diseases emerge from case studies, tracing human phenotypes to mutations of mtDNA (top-down). There are only few studies, however, taking the inverse approach, i.e. to follow up particular mutations to establish their impact on pathogenicity (bottom-up), although initial whole exome sequencing studies in patients have been undertaken which identified around 1% of mtDNA mutations in neurological patients [33] . Most bottom-up animal studies on mtDNA mutations, in turn, use the mtDNA mutator mouse model, expressing a proofreading deficient version of the nucleus-encoded catalytic subunit of mtDNA polymerase-*γ* (*PolgA*). This model accumulates increasing levels of somatic point mutations in mtDNA during its lifetime [34], somewhere in the mitochondrial genomes. In the current study, we take a different approach by studying conplastic mouse models, carrying stable point mutations in mitochondrial genes. Importantly, these models carry *defined*, *non-accumulating* mutations during their whole life, which is important, given that the rate of ageing and related phenotypes may be set early in life [35].

Since mtDNA mutations affecting either OXPHOS [36 – 40] or tRNA have been linked to ageing or neurological syndromes, we wanted to gauge the differential contribution of such mutations, and their combination, to ageing and neurological dysfunction. We were specifically interested in the influence of adenine insertions in the arm of the DHU-loop of mt-tRNA(Arg), since it was shown before that mutations of the tRNA body can generally affect translation, even located far apart from the anticodon [41]. Hence, three strains (one affecting OXPHOS alone, one affecting tRNA and one affecting both) (Table 2) were compared. To unveil the impact of these mutations, we correlated life-span data to performance in complex traits, such as learning and memory, and explored cellular mechanisms with respect to ageing and neurodegeneration (synaptic plasticity, reactive oxygen species (ROS), and shift in glia-neuron ratio).

## Results

### The lifespan of high-A + OXPHOS mutated mice is significantly shorter

As mitochondrial dysfunctions are heavily implicated in the ageing process [42], we were interested to explore the association of an adenine insertion in the mt-tRNA(Arg) (“high-A”) and an electron transport chain (“OXPHOS”) mutation with lifespan. A total of 42 mice (20 male, 22 female) carrying a combination of the high-A and OXPHOS mutation (Methods, table 2) were compared to 42 mice (21 per sex) with the OXPHOS mutation and 42 mice (21 per sex) with the high-A mutation. Survival curves for each strain are presented in Fig. 1. The median lifespan of high-A + OXPHOS mutated mice was significantly decreased by 7.11 % (60 days, from 843 to 783 days) compared to mice with OXPHOS mutation alone, and even more by 11.53 % (102 days, from 885 to 783 days) compared to mice with high-A mutation alone (Fig. 1; Table 1). This was independent of the sex of the affected high-A + OXPHOS mutation animals, as the difference of the survival values of males (mean: 791 days, 95% CI 723 - 859 days) and of females (mean: 768 days; 95% CI 651 -885 days) was not significant (p=0.417).

**Figure 1.**
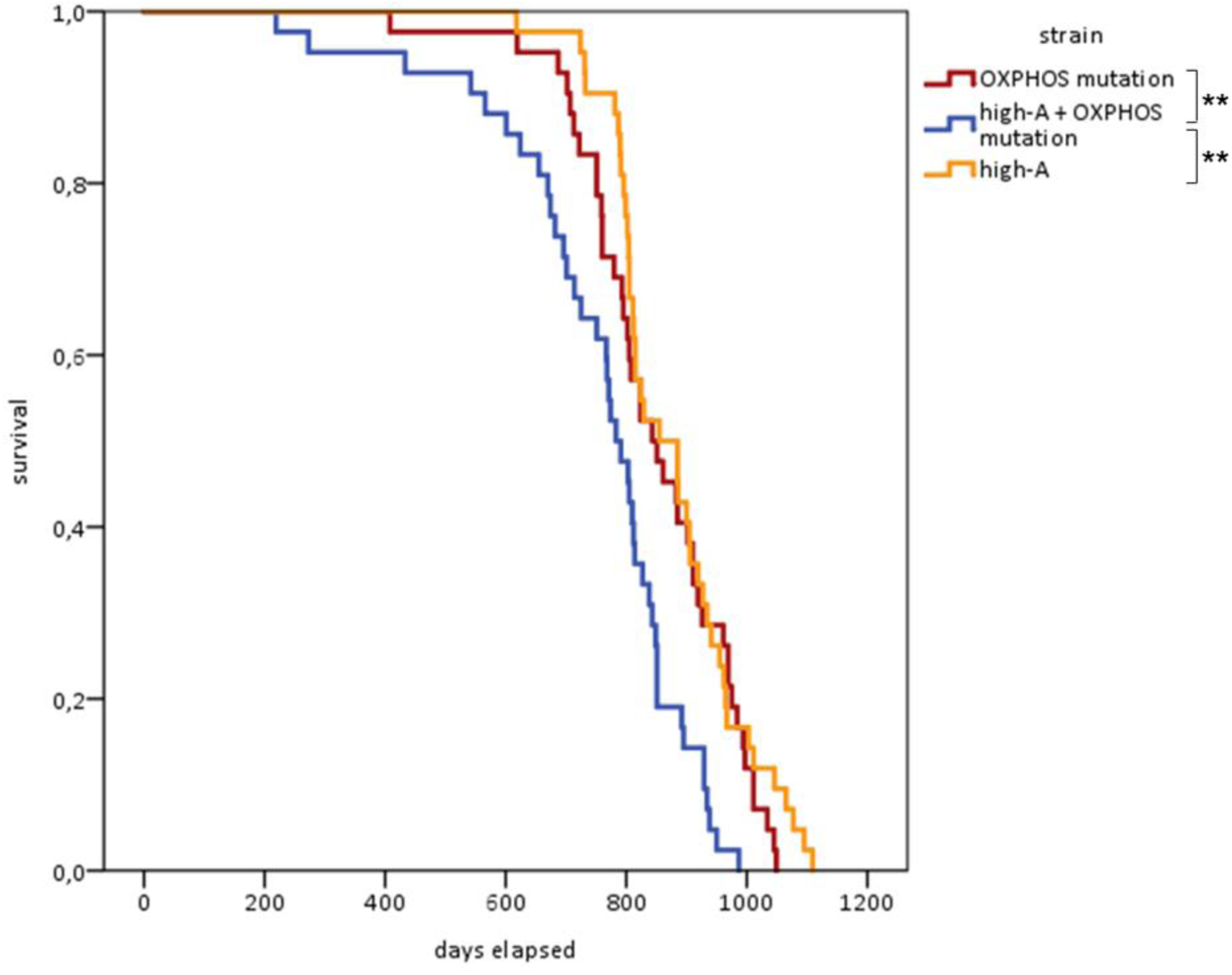
Survival analysis of high-A + OXPHOS mutated mice compared to mice that carry only one type mutation (OXPHOS or high-A alone). *Kaplan-Meier survival curves, Log-rank test, ** p < 0.01; OXPHOS mutation n = 42 (21 per sex) median survival 843 days; high-A + OXPHOS mutation n = 42 (20 male, 22 female) median survival 783 days; high-A n = 42 (21 per sex) median survival 885 days*.

**Table 1.**
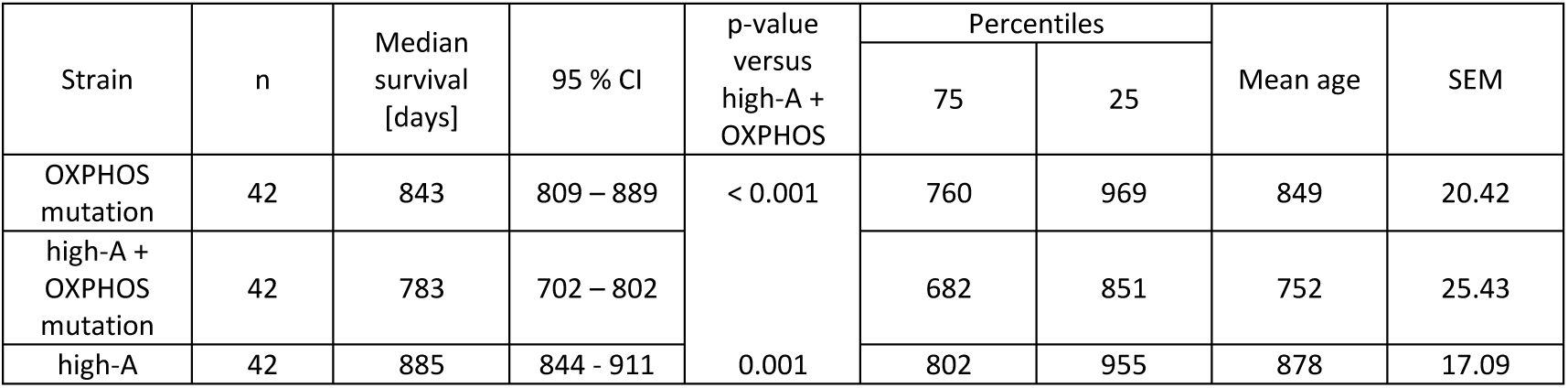
Lifespan analysis

### Aged high-A + OXPHOS mutated mice show poor physical performance in Morris Water Maze task

Since mitochondrial disorders may correlate with cognitive dysfunction [43], we hypothesized that the specific mtDNA mutations studied here have functional effects on higher brain performance. Therefore, we investigated the spatial learning ability of all conplastic mouse strains in Morris Water Maze (MWM) for three different age groups (3, 12 and 24 months) (Fig. 2). While 3- and 12-month-old mice did not show differences in MWM performance, 24-month-old high-A + OXPHOS mutated mice were regularly unable to complete the MWM task (i.e. stopped active swimming and were in danger of drowning), suggesting poor physical constitution in this strain and precluding cognitive performance comparison among the old age groups. Only one out of ten mice was able to complete all trials of the paradigm, while this rate was inversed and hence significantly higher (OXPHOS mutated mice 9 out of 10, high-A mice 8 out of 10) in strains with only one mutation (Fig. 2D) (p<0.01).

**Figure 2.**
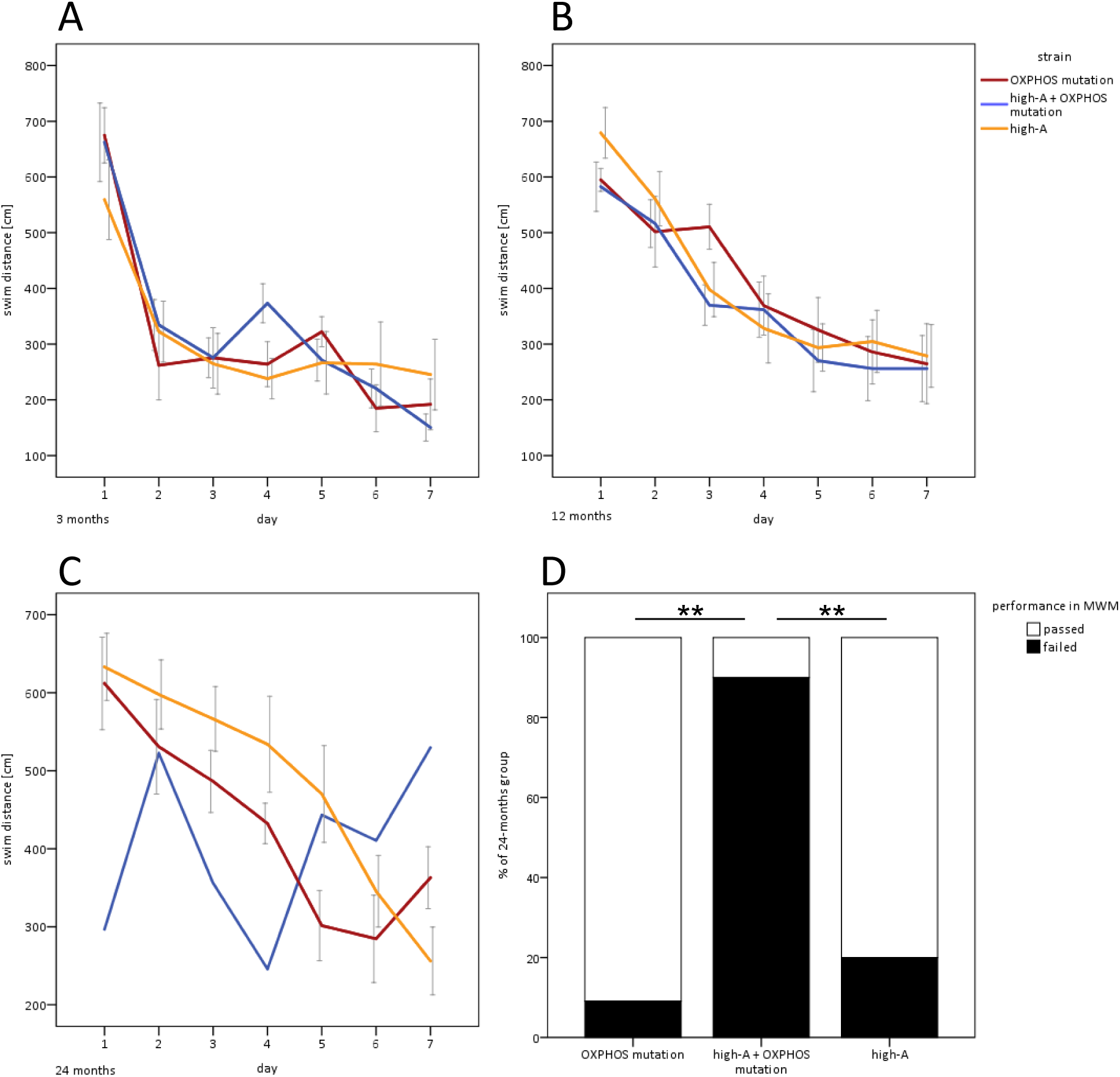
Aged high-A + OXPHOS mutated mice show impaired exercise ability in MWM task. *Morris Water Maze performance of 3-month (A), 12-month (B) and 24-month (C) old mice. OXPHOS mutated mice are shown in red (n = 9; 8; 10), high-A + OXPHOS mutated mice in blue (n = 10; 10; 1) and high-A mutated mice in orange (n = 10; 10; 8). A - C Learning curves of seven consecutive days (spatial acquisition). Data points represent mean values of swim distances of cohorts each day (± SEM). Note that in the high-A + OXPHOS group, only one animal was included, hence the blue curve does not show any SEM values. Kruskal-Wallis test followed by pairwise Mann-Whitney-U test. (D) Frequency distribution of animals that passed respectively failed in finishing Morris Water Maze task. From high-A + OXPHOS mutated mice only 1 out of 10 finished MWM. Chi-square test followed by pairwise Fisher’s exact test, ** p < 0.01*.

### No long-term potentiation (LTP) induction in young high-A + OXPHOS mutated mice

As we were not able to compare learning and memory differences in old mice using *in vivo* MWM, we resorted to *in vitro* methods, speculating that on the synaptic plasticity level, age-dependent changes might emerge. Hence, we investigated hippocampal long-term potentiation (LTP), which is thought to reflect learning mechanisms on the cellular network level. Surprisingly, LTP induced by theta-burst stimulation of CA1 pyramidal cells via Schaffer-collateral activation showed a difference between strains only in the youngest, 3-month-old group, but not in the adult or aged ones (12 or 24-month-old mice) (Fig. 3). Thus, while LTP could readily be induced in both single mutant strains (OXPHOS mutation 123 ± 9 %; high-A 132 ± 11 *%* ), there was virtually no LTP induction in 3-month-old high-A + OXPHOS mutated mice (100 ± 6 %). Hence, poor physical performance *in-vivo* late in life (perhaps masking possible cognitive performance decline) is preceded by synaptic-plasticity infringement on the cellular level, or possibly developmental plasticity delay, at early developmental stages.

**Figure 3.**
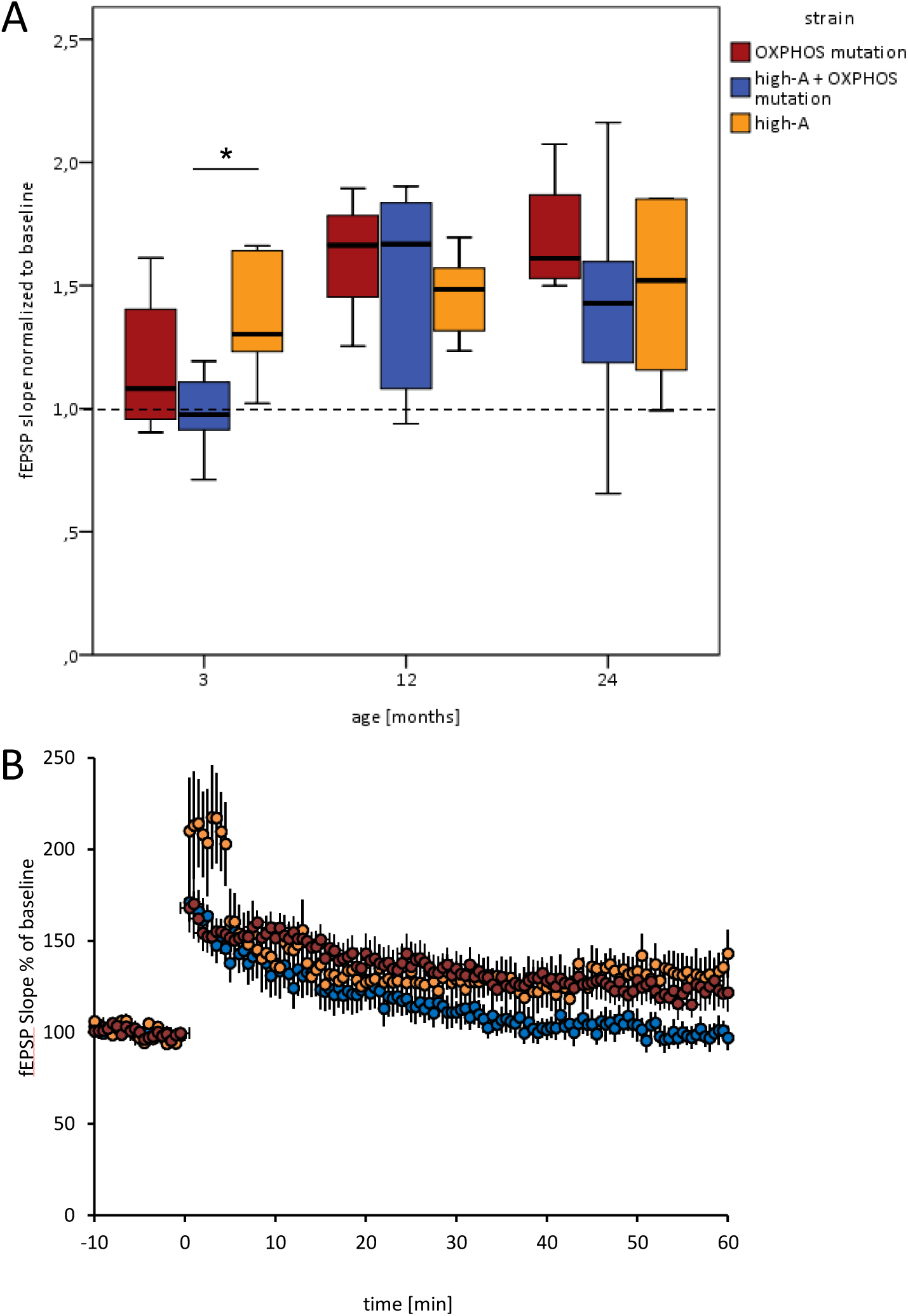
*No LTP induction in young high-A + OXPHOS mutated mice*. *(A) Box plots of LTP, calculated between minute 55 and minute 60 after TBS stimulation relative to baseline. 3-month-old high-A + OXPHOS mutated mice did not show LTP induction (100 ± 6% of baseline). (B) Time course of field excitatory postsynaptic potentials (fEPSP) measured in Schaffer collateral-CA1 synapses from 3 months old OXPHOS mutated (red, n = 13 slices /6 animals), high-A + OXPHOS mutated (blue, n = 10 slices / 7 animals), and high-A (orange, n = 11 slices /5 animals) mice. Each circle represents the percentage of fEPSP slope relative to mean baseline value. Following a 10 minutes baseline recording, three times of theta-burst stimulation protocol (TBS) was delivered at time point 0. Mann-Whitney-U test, * p < 0.05*.

### High-A, but not high-A + OXPHOS mutated mice show increased glial fibrillary acidic protein (GFAP) level at 24 months

Although hippocampal synaptic plasticity, i.e. LTP, was only affected early in life in high-A + OXPHOS mutated mice, we speculated that morphological changes might persist longer. Since activation of astrocytes with further astrogliosis is known to accompany many pathological conditions affecting the CNS [44], we focused on analysing this aspect. As astrocyte activation is characterized by increased expression of GFAP (glial fibrillary acidic protein), we quantified and then averaged the amount of GFAP in comparison to NeuN (neuronal nuclei) in the hippocampal cell layers CA1 (cornu ammonis area 1), CA3 (cornu ammonis area 3) and DG (dentate gyrus). In all three strains, the GFAP to NeuN ratio increased during ageing from 3 to 24 months (OXPHOS + 44.0 %; high-A + OXPHOS + 74.8 %; high-A + 153.5 %). While in both high-A strains the rise of GFAP signal occurred only late, i.e. at age 24 months, the strain with OXPHOS mutation alone showed an earlier, mid-life GFAP increase at 12 months (Fig. 4). Both strains with OXPHOS mutation only had moderate and insignificant elevations of GFAP signal, whereas the high-A strain showed a strong increase in GFAP to NeuN ratio (+ 153.5 %, p = 0.008).

**Figure 4.**
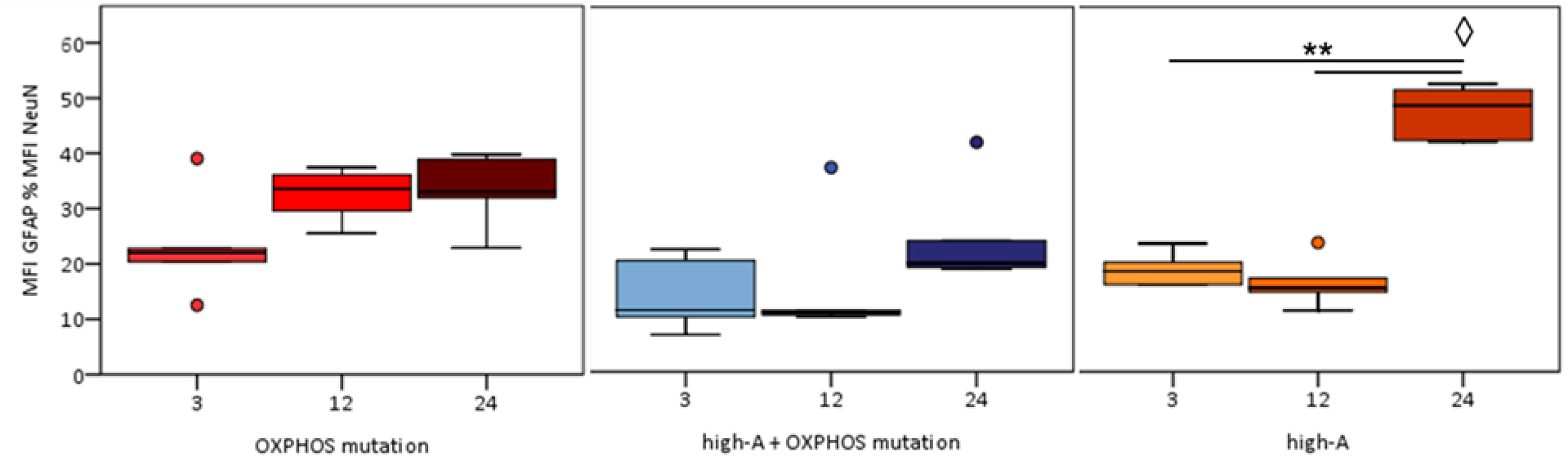
*GFAP to NeuN ratio in hippocampal cell layers*. *GFAP to NeuN ratio was measured by antibody staining of hippocampal slices. Box plots show comparison of GFAP to NeuN ratio in hippocampal cell layers from 3-, 12- and 24-month-old conplastic mutant strains (OXPHOS mutated shown in red, high-A + OXPHOS mutated in blue and high-A mice in orange). Each bar represents mean fluorescence intensity (MFI) of GFAP as percentage to MFI of NeuN. All data shown with n = 5 (5 animals per strain per age group, 2 slices per animal, 3 pictures per slice). Mann-Whitney-U test, ** p < 0.01, ◊ significant difference to both other strains in the same age group with p < 0.01*.

### Elevated neocortical superoxide level in aged high-A + OXPHOS mutated mice

We next hypothesized that mtDNA alterations, especially OXPHOS ones, could be associated with changes of ROS production. We therefore quantified mitochondrial superoxide by fluorescence microscopy of MitoSOX staining.

In the hippocampus, the structure associated with both spatial learning *in vivo* and LTP-changes, both OXPHOS mutated mice (+ 27.7 % lifetime change, p = 0.032) and high-A + OXPHOS mutated mice (+ 19.8 % lifetime change, p = 0.008), showed a significant rise of superoxide during life from 3 to 24 months, starting with relatively low levels. By contrast, in tissue from high-A mice (+ 6.0 % lifetime change), superoxide levels started out at highest levels in juvenile tissue, and afterwards did not change significantly during ageing. The difference between the age-dependent dynamics of superoxide levels, i.e. increase in the case of OXPHOS mutation and no lifetime change in the case of high-A only mutation, may corroborate an age-dependent impact of the OXPHOS mutation on superoxide production, or perhaps more likely, an age-related decline of initially highly active protective mechanisms (Fig. 5), as SOD2 levels in brains of aged OXPHOS-mutated mice were generally lower than in the high-A group (Fig. 7) (see below).

**Figure 5.**
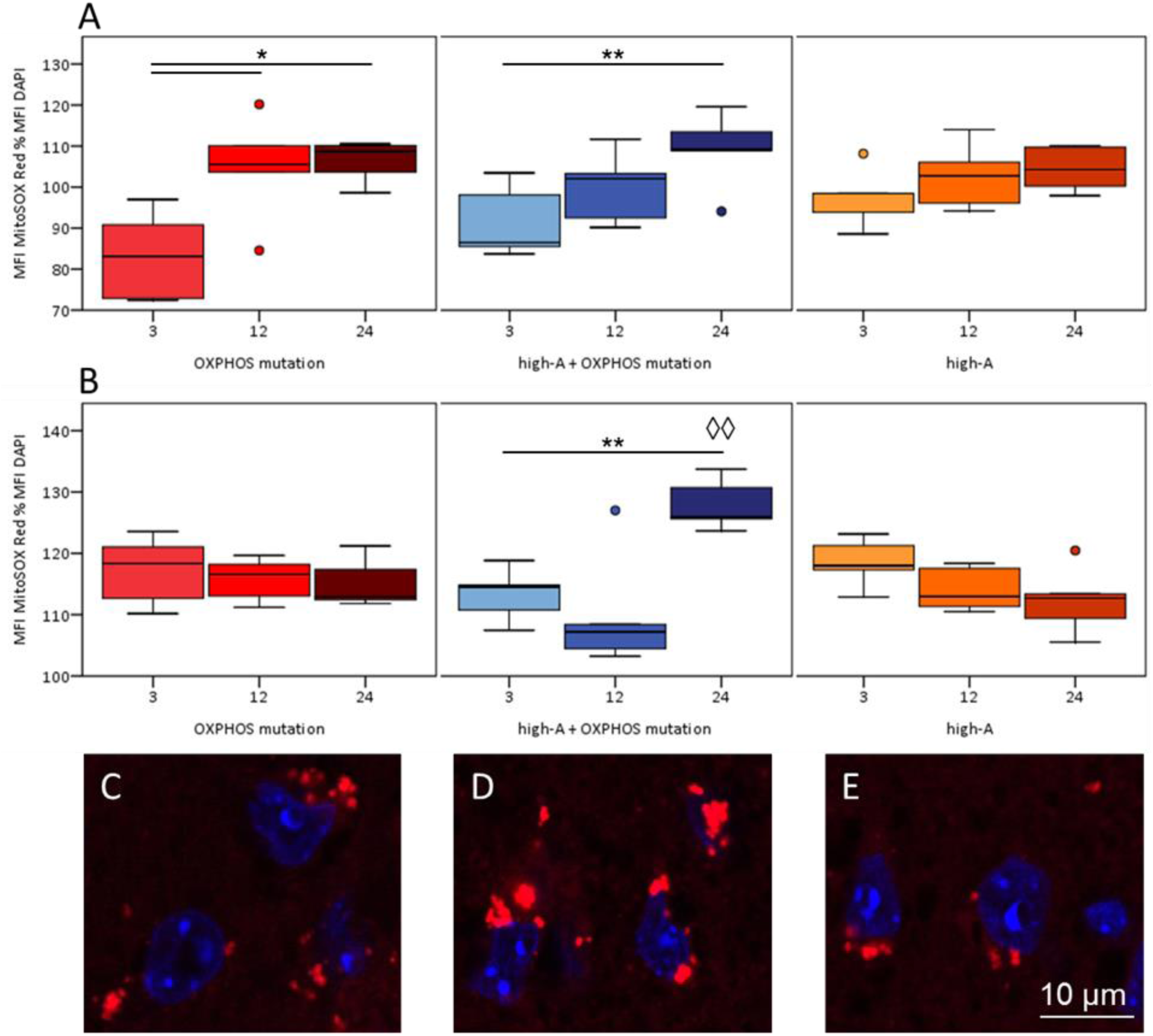
*Mitochondrial superoxide levels in hippocampus and neocortex*. *Mitochondrial superoxide production in hippocampal and neocortical slices was stained by MitoSOX Red solution (shown in red). Nuclei were counterstained with DAPI (shown in blue). Box plots illustrate comparison of mitochondrial superoxide levels in hippocampus (A) and neocortex (B) between conplastic mutant strains. Values represent mean fluorescence intensity (MFI) of MitoSOX Red as percentage to MFI of DAPI. All data shown with n = 5 (5 animals per strain per age group, 5 slices per animal, 6 pictures each slice). Mann-Whitney-U test, * p < 0.05, ** p < 0.01. (C-E) Sample pictures of neocortical cells from 24-month-old OXPHOS mutated (C), high-A + OXPHOS mutated (D) and high-A (E) mice. Pictures were taken with 120x magnification*.

We also analyzed mitochondrial superoxide in the neocortex, since besides possible hippocampal dysfunction, neocortical age-related morphological changes are known to correlate with low memory performance [45]. Importantly, only animals of high-A + OXPHOS group showed an increase (+ 13.3 %, p = 0.008) of superoxide during ageing (Fig. 5); compared to age-matched 24-month-old mice, the high-A + OXPHOS mutated strain had a significantly elevated neocortical superoxide level (+ 11.3 % compared to OXPHOS mutated, + 14.3 % compared to high-A, p=0.008), which might be linked to the low motor/physical performance in this group, as motor function is crucially dependent on neocortical function.

### OXPHOS mutations enhance mitochondrial network inhomogeneity in the hippocampus of aged mice

As increased ROS levels might have an impact on mitochondrial stability and network formation, we next analyzed mitochondrial morphology in 24-month-old mice with MitoTracker Deep Red. Abnormal mitochondrial network homogeneity can be detected by accumulation points of MitoTracker Deep Red fluorescence, which mark abnormal lumping of mitochondria [36]. We quantified pixels over threshold from MitoTracker fluorescence pictures (Fig. 6). Both OXPHOS mutated strains showed increased mitochondrial network inhomogeneity in the hippocampus of aged mice (+ 103 % OXPHOS mutated mice compared to high-A mice, p=0.032). Interestingly, both strains also show lower mitofusin 2 expression than the high-A strain (Fig. 7) (see below), which might explain this finding.

**Figure 6.**
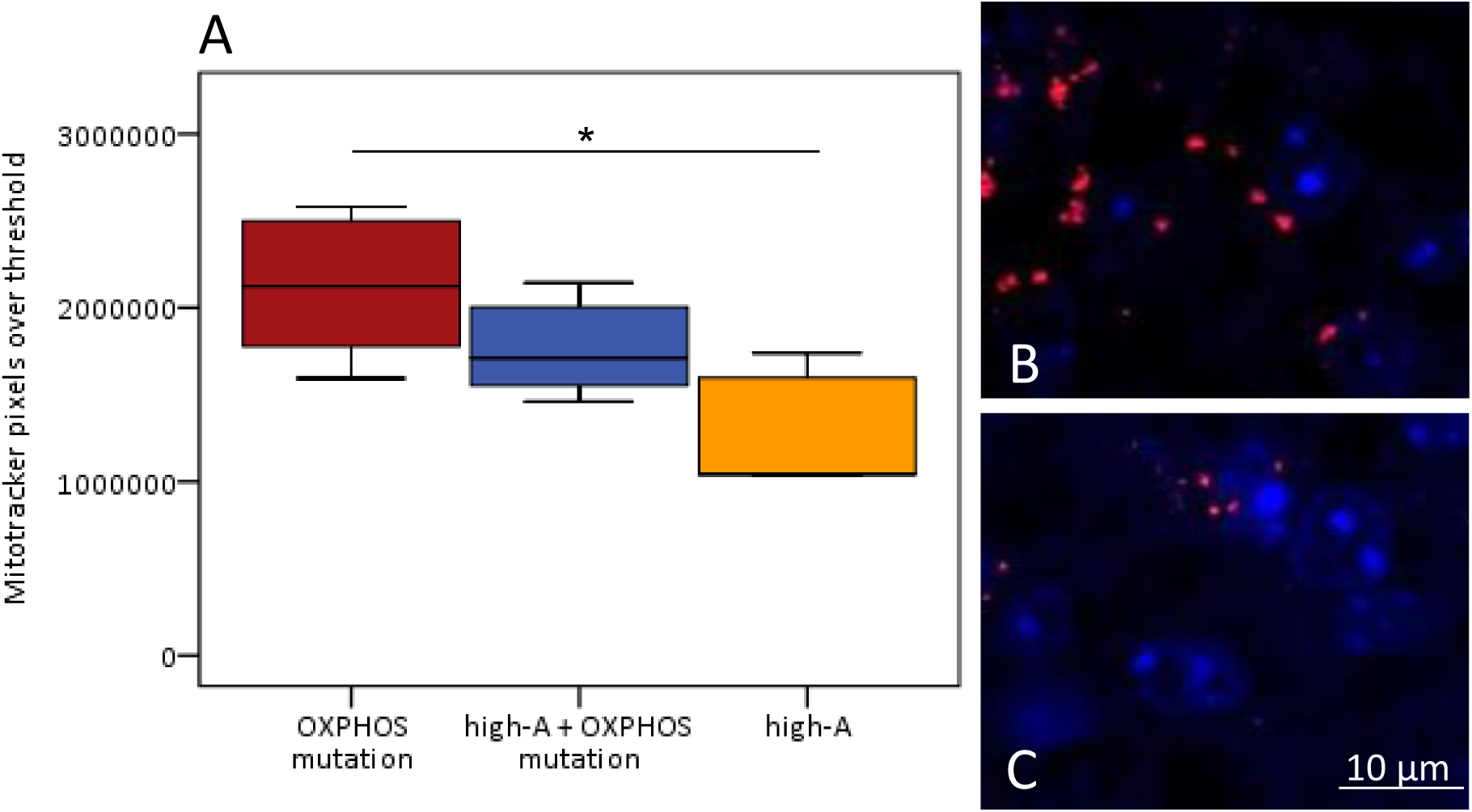
*Mitochondrial network homogeneity in the hippocampus of aged mice*. *(A) Box plot illustrates MitoTracker Deep Red fluorescence pixels over threshold. All data shown with n = 5 (5 animals per strain, 2 slices per animal, 3 pictures per slice). Mann-Whitney-U test, * p < 0.05 (B-C) Sample pictures of hippocampal cells from 24-month-old OXPHOS mutated (B), and high-A (C) mice. MitoTracker Deep Red pixels over threshold are shown in red, nuclei were counterstained with DAPI (shown in blue). Pictures were taken with 60x magnification*.

**Figure 7.**
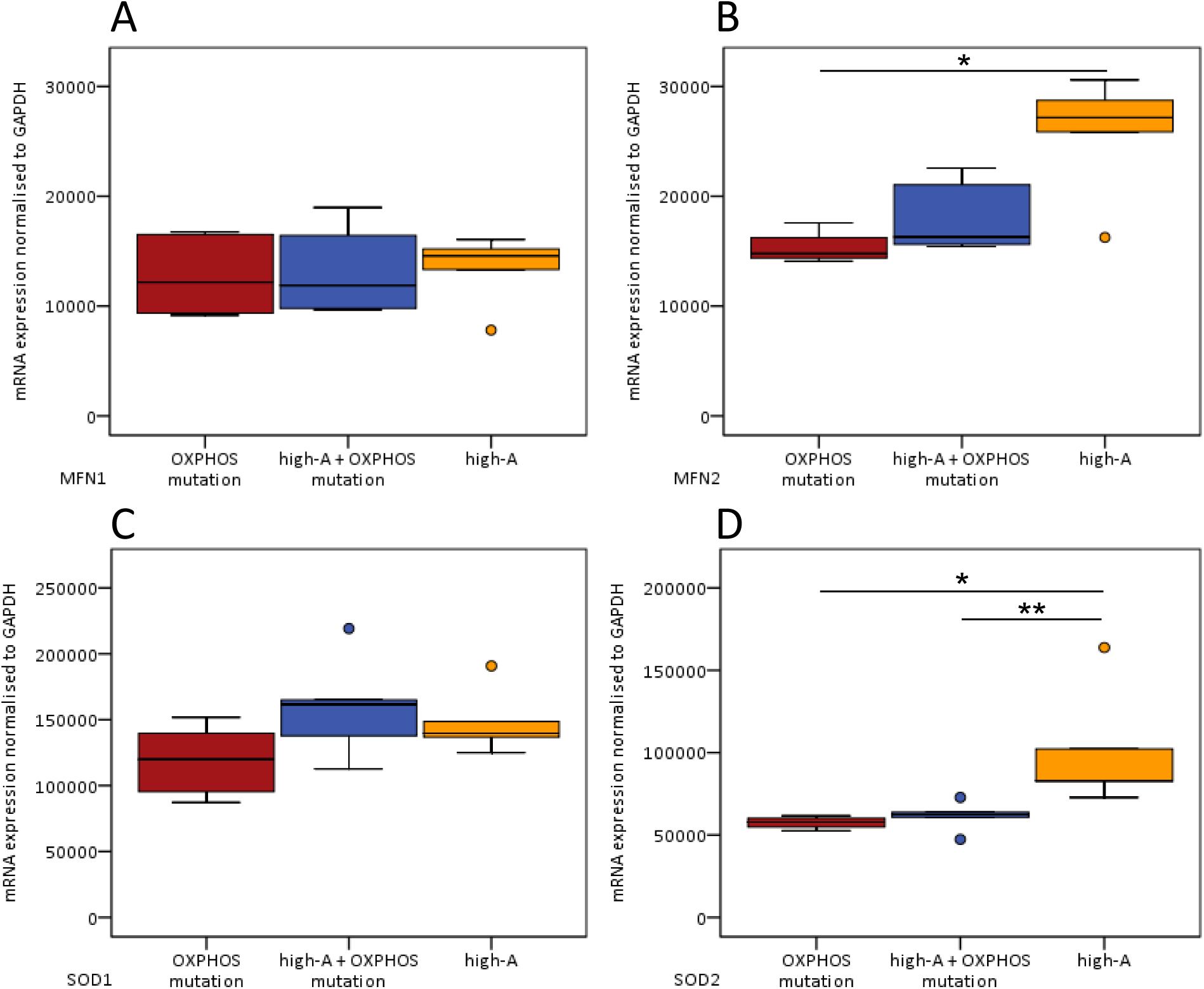
*Gene expression analysis from brain tissue of 24-month-old mice*. *Gene expression analysis of proteins pivotal in mitochondrial antioxidative response and mitochondrial network formation. Gene expression of the mitochondrial fusion contributor MFN2 (B), but not MFN1 (A) is lower in OXPHOS mutated mice. Gene expression of antioxidative enzyme SOD2 (D), but not SOD1 (C), is lower in OXPHOS mutated mice. All data shown n = 5. Mann-Whitney-U test, * p < 0.05, ** p < 0.01*.

There was no such difference in the neocortex (data not shown). This tissue specific difference perhaps indicates a special vulnerability of the hippocampus to metabolic stress, which under certain circumstances has been reported to be compensated for in the neocortex by protective antioxidant mechanisms, such as higher glutathione levels [46].

### OXPHOS mutated mice differ in MFN2 and SOD2 gene expression

To obtain more detailed information on further age-related mitochondrial changes, we analyzed the gene expression of proteins pivotal in mitochondrial metabolism, antioxidative response and mitochondrial network formation. In the brain of 24-month-old mice only two of gene expression values were significantly different (Fig. 7).

Expression of *Mitofusin 2* (MFN2, primarily expressed in brain), a contributor to mitochondrial fusion, was significantly lower in strains with OXPHOS mutation, while MFN1 (also present in brain, but primarily in other tissues) [47] expression was the same for all three strains. In OXPHOS mutated mice, MFN2 mRNA content was reduced to 60 %, in high-A + OXPHOS mutated mice even more to 54 % of the gene expression in high-A mice, corresponding to the findings on fusion-fission deficiencies shown above.

Expression of SOD2, the mitochondrial manganese superoxide dismutase, was significantly higher in the brain of high-A mice (133 % of OXPHOS mutated; 143 % of high-A + OXPHOS mutated), while SOD1, the cytosolic superoxide dismutase, was similarly expressed in all three strains. Hence, OXPHOS strains had lower mitochondrial SOD levels at old age, corresponding partially with the findings on superoxide levels shown above.

## Discussion

Large-scale mtDNA defects are widely known to have a negative impact on lifespan and tissue integrity. The present study asks the question whether also relatively isolated and specific point mutations in the mitochondrial genome can have observable impact on global life parameters such as lifespan and cognitive function, and whether a combination of two defects causes additive effects. On one hand, one can readily expect that mutations in genes of the mitochondrial respiratory chain responsible for oxidative phosphorylation (OXPHOS) can cause mitochondrial dysfunction [48]. On the other hand, mitochondrial tRNA alterations seem to be of particular importance for mitochondrial dysfunction, keeping in mind that genes coding for tRNAs represent only 10 % of the total mtDNA sequence, but are responsible for half of mtDNA mutational diseases [49]. Moreover, we examined the combination of an OXPHOS single nucleotide polymorphism (SNP) and a tRNA mutation to elucidate if there is a combination effect.

### Age gap between spatial learning and synaptic plasticity deficiency

While the OXPHOS only and mt-tRNA only mutation strains showed no or only moderate age-dependent changes in synaptic plasticity and superoxide and in particular reached a ceiling at mid-age in the latter parameter, the combined mutation showed clear plasticity defects and highest superoxide levels. We had expected to see such a correlation between spatial learning and *in vitro* parameters of hippocampal cell alterations, e.g. superoxide levels, with synaptic plasticity (measured as LTP) as the link in between. Puzzlingly, for the combined mutation this was the case in our study, but only with an age gap: at juvenile age, learning / physical condition was best, superoxide not elevated and synaptic plasticity, in turn, absent. In the old age group, synaptic plasticity normalized, while the other parameters deteriorated. How could this discrepancy be explained? The hippocampal formation has been identified for many years as a key structure in consolidation of spatial memory [50]. LTP, initially established *in vitro* by LØmo [51], was suggested to be one main cellular mechanism of learning [52] because this postsynaptic plasticity form shows stable enhancement of synaptic transmission after many hours [52, 53]. ROS was shown to play an important role in the expression but also the suppression of LTP [54]. Crucially, an increasing number of studies show that indeed the link between learning behavior and LTP is far from being direct. Thus, a review by Lynch [55] lists both, studies confirming correlations between cognitive ability and LTP, and others dismissing them. An example for such dissociation between learning performance and LTP was described by Huang *et al.* in 2013 [56] for a mouse model of ageing, where in fact reduced learning performance was associated with increased LTP. We suggest that synaptic plasticity dysfunction could act as an *early* marker of future higher-function decline.

### Superoxide levels, electron-transport-chain challenge, and poor physical condition

One *in vitro* parameter, for which high-A + OXPHOS mutated mice differed in our study, was neocortical superoxide which showed a strong increase in 24-month-old mice. How can this contribute to the observed phenotype of impaired ability to perform an exercise such as the water maze task? As some of the neocortical functions (mainly sensory perception, generation of motor commands and spatial reasoning) [57] are essential for the water maze task, changes in superoxide content might destabilize the underlying cellular mechanisms, as balanced superoxide levels are known to be important for cellular function [54, 58]. In addition to this, one cannot exclude that peripheral effects on other organ systems harboring the same mutation contributed to reduced exercise ability.

The other *in vitro* parameter, for which high-A + OXPHOS mutated mice differed, was the reduced MFN2 expression together with changes in mitochondrial network formation, as aged mice with the combined mutation tended to have enhanced network inhomogeneity in hippocampal cells. As this finding was seen in a similar way in mice with OXPHOS mutation alone, it is likely that electron transport chain (ETC) mutations in general predispose to this condition. This finding is in line with recent evidence that MFN2 not only mediates mitochondrial outer membrane fusion, but also has additional functions potentially required to sustain cellular energy demand [59]. Thus, a set of *Mfn2* knockout mice, developed by Chen *et al.* [60], are either lethal early postnatally (complete KO), or show severe tissue-specific changes with only partial loss of MFN2. In neurons, the loss of MFN2 is associated with a loss of complex IV activity [60] which indicates a link between the ETC and MFN2.

Lastly, we found higher GFAP levels together with higher SOD2 expression – but in this case most pronounced in the aged mice of high-A strain, and not or not as prominent, in both OXPHOS mutations strains. While it is a reactive sign, astrocyte activation seems to be important to maintain or retain brain integrity as much as possible in the acute phase of traumata and oxidative stress [44]. Thus, Thomas *et al.* [61] described a 3-fold increase of GFAP in the thalamus of rats after traumatic brain injury, and Daverey and Agrawal [62] showed astrocyte activation after mild oxidative stress. The latter study concluded the upregulation of GFAP to be an initial response mechanism of astrocytes to protect themselves under stress conditions. SOD2 is considered to be a mitochondrial superoxide scavenger, but also, and perhaps even more importantly, its major function is related to redox signalling mechanisms [63]. Therefore, the increased GFAP level in high-A mice might be a first sign, and the enhanced SOD2 expression a first defence mechanism, for a challenge to hippocampal function in these mice. However, without the additional OXPHOS mutation of the high-A + OXPHOS strain, the mt-tRNA mutations appear to be compensated for, and stay below threshold for a pathological phenotype on organism level.

### Combination effects of mutations

As our study shows, the combination of two point mutations, the mutation affecting OXPHOS (respiratory chain complex IV) and the adenine insertion in mt-tRNA(Arg), clearly reduces lifespan and exercise ability in the course of ageing; neither the strain with OXPHOS (complex V) mutation alone, nor the one with high-A content in mt-tRNA(Arg) alone show reduced lifespan or impaired exercise ability. But is it really due to combination of mutations or can we attribute the effect to only one of the players? Although mt-tRNA mutations are reported to be related to mitochondrial dysfunction associated exercise intolerance [28; 32], the mt-tRNA(Arg) alteration alone cannot be causal. Our results clearly show that the high-A strain does not share phenotypical similarities with the high-A + OXPHOS strain, at least in the parameters we analyzed. Indeed, in this strain, a number of (compensatory?) changes emerge, particularly in the oldest age group: Although the production of superoxide is similarly high in high-A and OXPHOS mutations (Figs. 5, 8), the scavenger SOD2 is upregulated in the high-A mice only. In a similar way, also mitofusin 2 is upregulated in this strain when compared to both OXPHOS mutations (Figs. 7, 8), which might explain relatively lower mitochondrial homogeneity, and hence more physiological fission-fusion behaviour. Does this mean this strain is completely unchallenged? As discussed above, the fact that there is increasing hippocampal astrocytosis and also an increase, albeit moderate, of hippocampal superoxide (similar to the OXPHOS-mutation alone; Fig. 8) suggests that the mutation does pose a challenge, but that this is levelling out at mid-age, perhaps due to the above-mentioned compensation.

**Figure 8.**
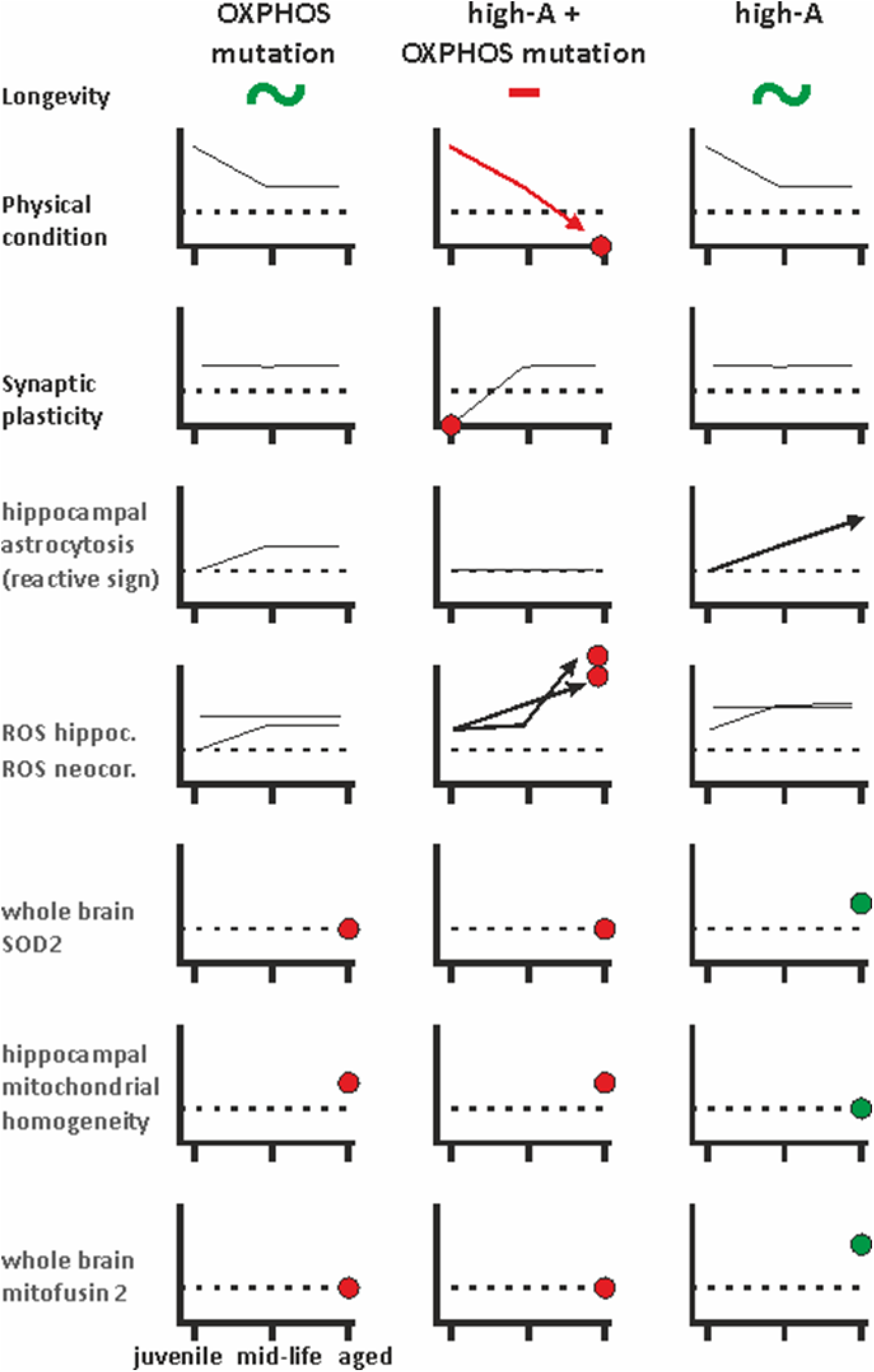
*Synoptic cartoon of age-dependent correlations*. *Functional readout parameters (black; longevity, physical condition, synaptic plasticity), and reactive (hippocampal astrocytosis) as well as mitochondrial-function-related changes (grey; superoxide levels, SOD2 expression, mitochondrial homogeneity, mitofusin 2 expression). Age groups are denoted on abscissa in lower left graph. Arrows show continuous age-dependent increases or decreases (no ceiling reached at mid-age). Red dots denote potentially detrimental and green dots potentially protective alterations*.

For the complex IV mutation the matter seems a bit different, and also not easy to clarify completely: As there is no existing mouse model carrying only the SNP at nt9348 leading to the alteration of Cytochrome c oxidase subunit III, a real control to this strain is lacking. Nevertheless, from other mouse models harboring a respiratory chain mutation, i.e. a complex IV deficiency, we know a wide range of different phenotypes [48]: Radford *et al.* developed a mouse model in which the nuclear encoded subunit CoxVIaH was knocked out [64]. CoxVIaH^-/-^ mice displayed a reduction in complex IV activity to about 23 % of the controls in cardiac tissue. Interestingly, as in our strain, these mice were viable and had normal lifespan. SURF1, in turn, is a protein known to participate in the early assembly steps of complex IV, and mutations in *surf1* cause severe neurological diseases in humans such as Leigh Syndrome, a disorder usually resulting in early lethality. *Surf1* knockout mice, generated by Dell'agnello *et al.* [65], showed mild complex IV deficiency (30 - 40 % of control values) in brain, heart, liver and skeletal muscle. Surprisingly these mice were phenotypically normal with even enhanced memory performance and a prolonged lifespan (5 months longer than control). *Sco2* encodes a chaperone that is required for assembly and function of complex IV. Whereas most SCO2 deficient humans die in infancy, Yang *et al.* [66] created two mouse models (*Sco2*^KI/KI^ and *Sco2*^KO/KI^) that showed complex IV deficiency in heart, muscle, brain and liver and also an accumulation of complex IV assembly intermediates in brain and liver, but no reduction of lifespan. One exception of a complex IV deficiency with reduced lifespan are the different types of *Cox10* knockout mice. The product of the *Cox10* gene catalyzes the first step of the biosynthesis of heme *a*, an essential prosthetic group for the function of complex IV. Diaz *et al.* developed a set of tissue-specific *Cox10* knockout mice, e.g. the *Cox10-Mlc-1f KO* (a myopathy model) [67]. These mice have a severe complex IV deficiency in skeletal muscle (13 % of residual activity) progressing with age and a very early death at about 6 months. Another exception, *Cox10-CaMKIIa* knockout mice, serve as an encephalomyopathy model because these animals develop a *progressive* complex IV deficiency which is restricted to forebrain structures [68]. They show behavioral abnormalities and severe cortical atrophy which results in premature death between 10 to 12 months of age. Neurons in the cingulate cortex and hippocampus are particularly affected by the mitochondrial defect. Gliosis and increased oxidative stress were observed in these parts of the brain. In a comparative view, our complex IV OXPHOS mutation strain shows a mild challenge (increasing superoxide up to mid-age) and comparatively lower levels of SOD2 and mitofusin 2 (Fig. 8).

Taken together, we conclude that it is the synergistic combination of both mutations (high-A mt-tRNA(Arg) + OXPHOS complex IV) which breaches a threshold and entails a shorter-lived phenotype with poor physical performance at old age. Potentially protective mechanisms that we saw to be triggered by the high-A mutation do not seem to enable the mice to cope with the additional challenge of the OXPHOS mutation so that eventually the tRNA mutation intensifies the phenotypic effects of OXPHOS mutation. For the OXPHOS mutation alone, it is likely that protective mechanisms exist as well, but that these were not reflected in the readouts measured in our study. Future studies could address the question whether the detrimental effect of the combined mutations on lifespan and exercise ability is due to an early onset, corresponding to early life effects as discussed by Ross et al. [35], or to a breakdown just in time with the occurrence of the phenotype at the age of 24 months, such as an accumulation over lifetime that destroys a system balance when a certain threshold is exceeded. In our case, the lack of LTP induction in 3-month-old high-A + OXPHOS mutation mice is hinting at an early onset.

## Methods

### Animal housing and care

All experiments were performed according to the guidelines of the local animal use and care committee, which also approved this study (Landesamt für Landwirtschaft, Lebensmittelsicherheit und Fischerei Mecklenburg-Vorpommern, permit number: 7221.3-1.1-059/12; Animal Care and Use Committee Lübeck, permit number: V242-7224. 122-5, Kiel, Germany). Animals were housed in groups up to 4 animals per cage. Cages were equipped with nesting material and red polycarbonate houses (i.e. environmentally enriched conditions). Animals were kept under stable surroundings (room temperature 23 ± 2 °C, relative humidity 40 ± 5 %, day-night rhythm with illumination 6 a.m. – 6 p.m.). Water and food were available *ad libitum*. For experiments, animals were taken from the housing unit in randomized fashion to reduce systematic bias.

### Conplastic mouse strains

Conplastic mouse strains of single mutations of mt-tRNA(Arg) or mitochondrial electron-transport-chain (ETC/OXPHOS) proteins were generated as described previously [69].

The C57BL/6J-mt^MRL/MpJ^ (referred to as *high-A*) strain carries a heteroplasmic stretch of adenine repetitions at mt.9821 in their mtDNA. As described by Sachadyn *et al.* [70] the poly(A) tract in the DHU arm of the tRNA(Arg) is heteroplasmic and varies from 9 to 14 A, where the prevailing variants are 10, 11 and 12 A. Checking for age-related changes by sequencing the mitochondrial genome revealed an additional heteroplasmic T to C mutation in mt.3900. This effects mitochondrial tRNA(Met) and was predicted to increase the length of the tRNA stem and decrease the size of the single stranded TΨC loop but not to change tRNA stability [70]. Heteroplasmy at this site ranged from 41 % to 87 %, with an average of 59 %, and did not change with age. As Chomyn et al. [71] described a 90% threshold level for tRNA mutations to cause respiratory chain dysfunction, it seems unlikely that this mutation contributes much to the phenotype of high-A strain.

The C57BL/6J-mt^FVB/NJ^ (referred to as *OXPHOS mutation*) harbors a SNP at position mt.7778 (7778 G>T) in the gene *mt-Atp8*, causing an Asp-Tyr amino acid exchange in the mitochondrial membrane ATP synthase subunit 8 in complex V.

The C57BL/6J-mt^NOD/LtJ^ (referred to as *high-A + OXPHOS mutation*) combines both, an ETC complex IV mutation and adenine insertion in the DHU-loop of tRNA-Arg. The SNP at mt.9348 (9348 G>A) in *mt-Co3* leads to an alteration of Cytochrome c oxidase subunit III (Val-Ile exchange) (Table 2).

**Table 2.**
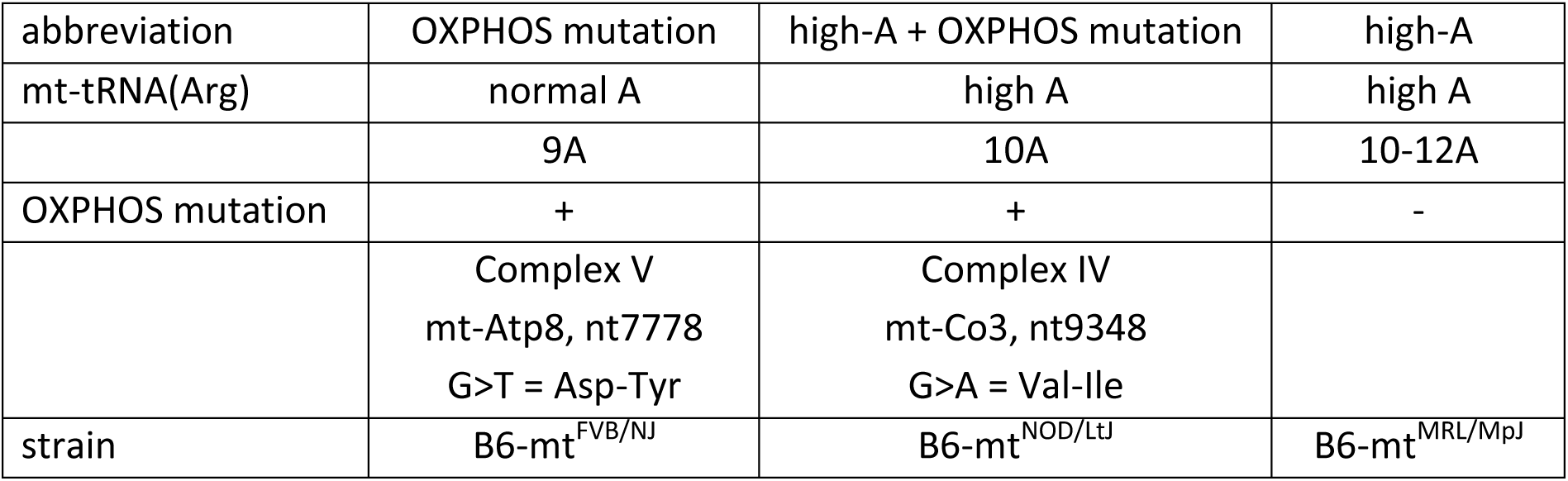
Conplastic mouse models used in this study

| abbreviation | OXPHOS mutation | high-A + OXPHOS mutation | high-A |
| --- | --- | --- | --- |
| mt-tRNA(Arg) | normal A | high A | high A |
|  | 9A | 10A | 10-12A |
| OXPHOS mutation | + | + | - |
|  | Complex V<br>mt-Atp8, nt7778<br>G>T = Asp-Tyr | Complex IV<br>mt-Co3, nt9348<br>G>A = Val-Ile |  |
| strain | B6-mt<sup>FVB/NJ</sup> | B6-mt<sup>NOD/LtJ</sup> | B6-mt<sup>MRL/MpJ</sup> |

### Lifespan study design

Fourty-two mice per group (OXPHOS mutation: 21 each sex; high-A + OXPHOS mutation: 20 male, 22 female; high-A: 21 each sex) were used for a longitudinal study to evaluate lifespan. All of them experienced a Morris Water Maze task once in their lifetime.

Animals were housed as described above and periodically inspected by certified personnel until natural death. A mouse showing more than one of the following clinical signs was determined moribund [40]: inability to eat or drink; severe lethargy (reluctance to move when gently prodded with forceps); severe balance or gait disturbance; rapid weight loss; an ulcerated or bleeding tumor; enlarged abdomen. Moribund mice were killed and the age was taken as the best available estimate of its natural lifespan.

### Spatial learning (Morris Water Maze)

Assessment of spatial learning abilities as a paradigm of cognitive function took place in an open-field variant of the Morris Water Maze (MWM) as described before [72]. Testing occurred in a separate room with stable temperature (air: 22 ± 1 °C, water: 21 ± 1 °C), brightness (110 lux) and minimal noise. For habituation mice were housed in the testing room 2 days prior to MWM experiments. Mice were generally healthy and free of open wounds. As visual ability is essential for spatial learning in the MWM, mice were checked for sight disorders [73] and if so excluded.

For MWM experiments visible cues (black geometrical symbols; 0.5 × 0.5 m) were attached to each wall. Mice had to reach a hidden platform (ø 7.5 cm) to escape the tank (ø 1.1 m) filled with opaque water. A submerged platform (target), approximately 1 cm below the surface of water was placed in a specific quadrant. Mice were randomly released from eight starting locations. After first day of habituation each mouse was tested six trials per day, seven consecutive days. Each trial had a time limit of 60 s, followed by 30 s remaining upon the platform and resting in a neutral cage for 60 s. Mice were gently guided to platform by hand, in case of failing to reach the target within time limit. Testing started at 09:00 a.m.each day. Tracks were recorded with a camera for subsequent data analysis with the software Etho Vision 3.1 (Noldus, Netherlands).

### Preparation of brain slices for electrophysiology and staining

Mice were deeply anesthetized via diethyl ether inhalation (Mallinckrodt Baker, Deventer, Netherlands) and thereupon decapitated. The brain was quickly removed and transferred into chilled and oxygenated (carbogen 95% O_2_ / 5% CO_2_) dissection solution (87 mM NaCl, 25 mM NaHCO_3_, 2.5 mM KCl, 1.25 mM NaH_2_PO_4_, 0.5 mM CaCl_2_, 7 mM MgCl_2_, 10 mM D-glucose, 75 mM sucrose adjusted to pH 7.4 and 326-328 mosmol/l H_2_O). The brain was glued to a vibratome (Integraslice 7550 MM, Campden Instruments Ltd., England) and slices were prepared in chilled and oxygenated artificial cerebrospinal fluid (aCSF) - plane: transversal for hippocampal formation (400 μm) and coronal for neocortex (500 μm) respectively. ACSF was comprised of 124 mM NaCl, 26 mM NaHCO_3_, 3 mM KCl, 1.25 mM NaH_2_PO_4_, 2.5 mM CaCl_2_, 1.5 mM MgCl_2_ and 10 mM D-glucose adjusted to pH 7.4 and 304-312 mosmol/l H_2_O. After preparation, slices were transferred into a submerged-type aCSF-storage chamber for maintenance and kept for 1.5 hours, before starting recordings and staining procedures.

### Field potential recordings for analysis of synaptic plasticity

Evoked field potential recordings were performed in an interface-type recording chamber (BSC-BU, Harvard Apparatus Inc., U.S.A.). During recording, slices were perfused with prewarmed (Haake C10, Electron Corporation GmbH, Germany) and oxygenated (carbogen, 95% O_2_ / 5% CO_2_) aCSF with a continual flow of 2-3 ml/min (Perimax, Spetec GmbH, Germany). The temperature of the fluid in the recording chamber was kept constant at 32 ± 1°C (TC-10, npi electronic GmbH, Germany). Schaffer collaterals were stimulated with a bipolar platinum electrode (PT-2T, Science Products GmbH, Germany). Stimulation was controlled by a Master-8 pulse stimulator (A.M.P.I., Jerusalem, Israel) connected to a stimulus isolator (A365, WPI Inc., U.S.A.), applying a paired-pulse protocol with 40 ms inter-pulse interval (IPI) and an inter-stimulus interval (ISI) of 30 s (0.033 Hz). Baseline stimulation strength was determined generating input-output curves until saturation of amplitude of field excitatory postsynaptic potentials (fEPSP); for further stimuli, intensity was reduced to half-maximum intensity. Following 10 minutes of stable baseline recording, theta-burst stimulation (TBS; 3 trains 20 s apart; train of 10 epochs at 5 Hz containing 5 pulses each; duration 150 |as at 100 Hz) was used to evoke long-term-potentiation. Field EPSPs were recorded in stratum radiatum of the CA1 subfield using borosilicate glass pipettes (GB150-8P Science Products GmbH, Hofheim am Taunus, Germany) with a tip resistance of 2-3 MQ (pulled with PIP5 puller from HEKA Elektronik, Lambrecht, Germany), filled with aCSF containing an Ag/AgCl wire. Evoked fEPSPs were amplified and filtered at 1 kHz (EXT-10-2F, npi electronic GmbH). Recordings were digitized (Micro 1401mkII, CED Ltd., Cambridge, UK) and analyzed using Signal 2.16 (CED Ltd). Slope of fEPSPs were measured, displayed and plotted relative to mean of baseline.

### Quantification of mitochondrial superoxide

Acute brain slices were incubated in 1 μM MitoSOX Red (Life Technologies, Darmstadt, Germany) in oxygenated aCSF for 15 minutes at room temperature and protected from light. After washing for 5 minutes in aCSF, slices were directly fixed in 3.7% paraformadehyde, cryo-protected with 30% sucrose in 1 x phosphate buffered saline (PBS) overnight and frozen. Probes were cut into 10 μm slices, counterstained and mounted with ProLong Gold Antifade Reagent containing DAPI (Life Technologies). Quantifying analysis of ROS levels relative to nucleic area stained with DAPI were performed by confocal laser scanning microscopy (Fluoview FV10i, Olympus). Values are composed of samples from five animals per strain. Five slices of each animal with six pictures each (two pictures from three different regions) were analyzed. In hippocampal formation pictures were taken in cell layers CA1 (cornu ammonis area 1), CA3 (cornu ammonis area 3) and DG (dentate gyrus). In neocortical slices cell layers I / II, III / IV and V/VI were analyzed.

### Quantification of mitochondrial network structure

Acute brain slices were handled as described above. Probes were cut into 10 μm slices, stained with MitoTracker Deep Red (Molecular Probes) in 1:20000 dilution for 30 min at room temperature and protected from light. Slices were counterstained and mounted with ProLong Gold Antifade Reagent containing DAPI (Life Technologies). Quantifying analysis was performed by confocal laser scanning microscopy (Fluoview FV10i, Olympus). The inhomogeneity of the mitochondrial network structure was determined as the amount of pixels with a higher intensity than a defined threshold value using ImageJ 1.51k quantification software. Values are composed of samples from five animals per strain. Two slices per animal with three pictures each were analyzed.

### Neuronal cell composition (GFAP/NeuN-ratio)

Hippocampal slices were fixed in 3.7% paraformaldehyde, cryo-protected with 30% sucrose in 1 x P B S at 4 ° C overnight and frozen. Probes were cut into 20 |am slices, blocked with 10% bovine serum albumin, 0.05% Triton X-100 for 1 h at room temperature and incubated with rabbit polyclonal anti-NeuN antibody (1:400; 1 h at 37 °C; Abcam (Cambridge, UK)). Sections were washed three times in P B S for 15 min and exposed to mouse monoclonal anti-GFAP antibody (1:400; 1 h at 37 °C; Abcam). Washing was followed by application of secondary fluorescent-labeled antibodies Cy3 goat anti-rabbit IgG and Cy5 goat anti-mouse IgG (1:2000; 1 h at 37 °C; Thermo Fisher Scientific, Waltham, USA). Quantification was conducted with fluorescence microscopy (Leica DMI 6000B (Wetzlar, Germany)). Values are composed of samples from five animals per strain. Two slices of each animal with three pictures each were analyzed.

### Gene expression analysis

Gene expression analysis was done as described before [36]. 30mg of brain were homogenized in Lysing Matrix D tubes with a Fast Prep^TM^ homogenizer (MP Biomedicals, Solon, USA). RNA was isolated and purified using an RNeasy^®^ Mini Kit (Qiagen, Hilden, Germany). The Maxima™ First Strand cDNA synthesis kit for RT-qPCR (Thermo Scientific, Darmstadt, Germany) was used to synthesize cDNA. For real-time PCR, cDNA solutions containing TaqMan Universal Master Mix (Applied Biosystems, Darmstadt, Germany) and the corresponding TaqMan^®^ Gene Expression Assay (Applied Biosystems) of primer and gene probe (Table 3) were amplified and detected using a PikoReal Real-Time PCR System (Thermo Scientific). Glyceraldehyde 3-phosphate dehydrogenase (GAPDH) served as a housekeeping gene for nuclear-encoded genes. Gene expression values were calculated from the Cq values in the Thermo Scientific PikoReal Software 2.1 (Thermo Scientific).

**Table 3.** Quantitative real-time PCR probes (Applied Biosystems).

| Gene | Assay |
| --- | --- |
| CAT | Mm01340247_m1 |
| DJ1 | Mm00498538_m1 |
| DNM1L | Mm01342903_m1 |
| FIS1 | Mm00481580_m1 |
| GAPDH | Mm99999915_g1 |
| GPX1 | Mm00656767_g1 |
| MFN1 | Mm01289369_m1 |
| MFN2 | Mm01255785_m1 |
| OPA1 | Mm01349716_m1 |
| PINK1 | Mm00550827_m1 |
| PRDX3 | Mm00545848_m1 |
| SIRT3 | Mm00452131_m1 |
| SOD1 | Mm01700393_g1 |
| SOD2 | Mm00690588_m1 |
| TFAM | Mm00447485_m1 |
| UCP2 | Mm00627599_m1 |

### Statistical analysis

Statistical analysis was performed with IBM SPSS Statistics 24. Significance levels of p < 0.05 (significant) and p < 0.01 (highly significant) were used to evaluate the null hypothesis. Survival curves are shown as Kaplan-Meier plot. Analysis was done by log rank statistic followed by posthoc test. For analysis of MWM acquisition phase, a Kruskal-Wallis test followed by pairwise Mann-Whitney-U test was used. The frequency distribution of animals that passed respectively failed in finishing Morris Water Maze task was analyzed by chi-square test followed by pairwise Fisher’s exact test. Comparisons of independent groups were performed with Mann-Whitney-U test. Results are shown as Box-and-Whisker plots.

## Data availability

Data will be available upon publication as Excel files on the server of the Institute of Physiology under the following address: https://physiologie.med.uni-rostock.de/publikationen/publikationen-in-peer-review-journalen/

## Acknowledgements

This study was supported by BMBF grant ROSAge 31P6662, and FORUN programme funding no. 889062. GF has received funding from the European Union’s Horizon 2020 research and innovation programme under Grant agreement No 633589.We thank M.Sc. Anne-Marie Neumann for technical support. Furthermore, we thank Mrs. Hanka Schmidt for support in animal care.

## Author Contributions Statement

RK designed the experiments, overall design of collaborative project ROSAge was carried out by GF, SI and RK. GR and RK wrote the main text of the manuscript, with contributions by GF. All figures were prepared by GR. Experiments on animal behavior were done by JM, extracellular LTP measurements were carried out by TT and JM. Mitochondrial ROS analysis, MitoTracker deep Red analysis and GFAP to NeuN ratio measurements were performed by GR. Gene expression analysis was done by CJ. All authors contributed to interpretation of data; all authors reviewed the manuscript.

## Additional Information

*Competing financial interests*

The authors declare no competing financial interests.

## References

c1. Duchen MR. Contributions of mitochondria to animal physiology: from homeostatic sensor to calcium signalling and cell death. J. Physiol. 516, 1–17 (1999).

c2. Desagher S, Martinou J-C. Mitochondria as the central control point of apoptosis. Trends Cell Biol. 10, 369–377 (2000).

c3. Nunnari J, Suomalainen A. Mitochondria: In Sickness and in Health. Cell 148, 1145–1159 (2012).

c4. Zhang F, Broughton RE. Mitochondrial-Nuclear Interactions: Compensatory Evolution or Variable Functional Constraint among Vertebrate Oxidative Phosphorylation Genes? Genome Biol. Evol. 5, 1781–1791 (2013).

c5. Weinberg SE, Sena LA, Chandel NS. Mitochondria in the Regulation of Innate and Adaptive Immunity. Immunity 42, 406–417 (2015).

c6. Sun T, Qiao H, Pan P-Y, Chen Y, Sheng ZH. Motile Axonal Mitochondria Contribute to the Variability of Presynaptic Strength. Cell Rep. 4, 413–419 (2013).

c7. Lagouge M, Larsson NG. The role of mitochondrial DNA mutations and free radicals in disease and ageing. Journal of Internal Medicine 273, 529–543 (2013).

c8. Wallace, D. C. et al. Mitochondrial DNA mutation associated with Leber’s hereditary optic neuropathy. Science 242, 1427 (1988).

c9. Holt IJ, Harding AE, Morgan-Hughes JA. Deletions of muscle mitochondrial DNA in patients with mitochondrial myopathies. Nature 331, 717–719 (1988).

c10. Poulton J, , Macaulay V, Hennings S, Mitchell J, Wareham NJ. Type 2 diabetes is associated with a common mitochondrial variant: evidence from a population-based case-control study. Hum. Mol. Genet. 11, 1581–1583 (2002).

c11. Cortopassi GA, Shibata D, Soong NW, Arnheim NA. Pattern of accumulation of a somatic deletion of mitochondrial-DNA in aging human tissues. Proc. Natl. Acad. Sci. U.S.A. 89, 7370–7374 (1992).

c12. Larsson NG. Somatic mitochondrial DNA mutations in mammalian aging. Annual Reviews Biochemistry 79, 683–706 (2010).

c13. Gorman GS, Schaefer AM, Ng Y, Gomez N, Blakely EL, Alston CL, Feeney C, Horvath R, Yu-Wai-Man P, Chinnery PF, Taylor RW, Turnbull DM, McFarland R. Prevalence of nuclear and mitochondrial DNA mutations related to adult mitochondrial disease. Ann. Neurol. 77, 753–759 (2015).

c14. Lin MT, Beal MF. Mitochondrial dysfunction and oxidative stress in neurodegenerative diseases. Nature 443, 787–795 (2006).

c15. Kraytsberg Y, Kudryavtseva E, McKee AC, Geula C, Kowall NW, Khrapko K. Mitochondrial DNA deletions are abundant and cause functional impairment in aged human substantia nigra neurons. Nat. Genet. 38, 518–520 (2006).

c16. Sterky FH, Lee S, Wibom R, Olson L, Larsson N-G. Impaired mitochondrial transport and Parkin-independent degeneration of respiratory chain-deficient dopamine neurons in vivo. Proc. Natl. Acad. Sci. 108, 12937–12942 (2011).

c17. Oka S, Leon J, Sakumi K, Ide T, Kang D, LaFerla FM, Nakabeppu Y. Human mitochondrial transcriptional factor A breaks the mitochondria-mediated vicious cycle in Alzheimer’s disease. Sci. Rep. 6: 37889; DOI:10.1038/srep37889 (2016).

c18. Chang Y, Kong Q, Shan X, Tian G, Ilieva H, Cleveland DW, Rothstein JD, Borchelt DR, Wong PC, Lin CL. Messenger RNA Oxidation Occurs Early in Disease Pathogenesis and Promotes Motor Neuron Degeneration in ALS. Plos One 3(8), e2849 (2008).

c19. Federico A, Cardaioli E, Da Pozzo P, Formichi P, Gallus GN, Radi E. Mitochondria, oxidative stress and neurodegeneration. J. Neurol. Sci. 322, 254–262 (2012).

c20. Burté F, Carelli V, Chinnery PF, Yu-Wai-Man P. Disturbed mitochondrial dynamics and neurodegenerative disorders. Nat. Rev. Neurol. 11, 11–24 (2015).

c21. Twig G, Elorza A, Molina AJ, Mohamed H, Wikstrom JD, Walzer G, Stiles L, Haigh SE, Katz S, Las G, Alroy J, Wu M, Py BF, Yuan J, Deeney JT, Corkey BE, Shirihai OS. Fission and selective fusion govern mitochondrial segregation and elimination by autophagy. EMBO J. 27, 433–446 (2008).

c22. Knott AB, Perkins G, Schwarzenbacher R, Bossy-Wetzel E. Mitochondrial fragmentation in neurodegeneration. Nature Rev. Neurosc. 9, 505–518 (2008).

c23. Picard M, Wallace DC, Burelle Y. The rise of mitochondria in medicine. Mitochondrion 30, 105–116 (2016).

c24. Grady JP, Campbell G, Ratnaike T, Blakely EL, Falkous G, Nesbitt V, Schaefer AM, McNally RJ, Gorman GS, Taylor RW, Turnbull DM, McFarland R. Disease progression in patients with single, large-scale mitochondrial DNA deletions. Brain 137, 323–334 (2014).

c25. Parikh S, Goldstein A, Koenig MK, Scaglia F, Enns GM, Saneto R, Anselm I, Cohen BH, Falk MJ, Greene C, Gropman AL, Haas R, Hirano M, Morgan P, Sims K, Tarnopolsky M, Van Hove JL, Wolfe L, DiMauro S. Diagnosis and management of mitochondrial disease: a consensus statement from the Mitochondrial Medicine Society. Genet. Med. 17, 689–701 (2015).

c26. Picard M, Zhang J, Hancock S, Derbeneva O, Golhar R, Golik P, O'Hearn S, Levy S, Potluri P, Lvova M, Davila A, Lin CS, Perin JC, Rappaport EF, Hakonarson H, Trounce IA, Procaccio V, Wallace DC. Progressive increase in mtDNA 3243A>G heteroplasmy causes abrupt transcriptional reprogramming. Proc. Natl. Acad. Sci. 111, E4033–E4042 (2014).

c27. Schon EA, DiMauro S, Hirano M. Human mitochondrial DNA: roles of inherited and somatic mutations. Nat. Rev. Genet. 13, 878–890 (2012).

c28. Yarham JW, Elson JL, Blakely EL, McFarland R, Taylor RW. Mitochondrial tRNA mutations and disease. WIREs RNA 1, 304–324 (2010).

c29. Lehmann D, Schubert K, Joshi PR, Baty K, Blakely EL, Zierz S, Taylor RW, Deschauer M. A novel m.7539C>T point mutation in the mt-tRNAAsp gene associated with multisystemic mitochondrial disease. Neuromuscul. Disord. 25, 81–84 (2015).

c30. Roos S, Darin N, Kollberg G, Andersson GrÖnlund M, Tulinius M, Holme E, Moslemi AR, Oldfors A. A novel mitochondrial tRNA Arg mutation resulting in an anticodon swap in a patient with mitochondrial encephalomyopathy. Eur. J. Hum. Genet. 21, 571–573 (2013).

c31. Uusimaa J, Finnilä S, Remes AM, Rantala H, Vainionpää L, Hassinen IE, Majamaa K. Molecular epidemiology of childhood mitochondrial encephalomyopathies in a Finnish population: Sequence analysis of entire mtDNA of 17 children reveals heteroplasmic mutations in tRNA(Arg), tRNA(Glu), and tRNA(Leu(UUR)) genes. Pediatrics 114, 443–450 (2004).

c32. Blakely EL, Swalwell H, Petty RKH, McFarland R, Turnbull DM, Taylor RW. Sporadic myopathy and exercise intolerance associated with the mitochondrial 8328G>A tRNA^Lys^ mutation. J Neurol 254, 1283–1285 (2007).

c33. Yang Y, Muzny DM, Xia F, Niu Z, Person R, Ding Y, Ward P, Braxton A, Wang M, Buhay C, Veeraraghavan N, Hawes A, Chiang T, Leduc M, Beuten J, Zhang J, He W, Scull J, Willis A, Landsverk M, Craigen WJ, Bekheirnia MR, Stray-Pedersen A, Liu P, Wen S, Alcaraz W, Cui H, Walkiewicz M, Reid J, Bainbridge M, Patel A, Boerwinkle E, Beaudet AL, Lupski JR, Plon SE, Gibbs RA, Eng CM. Molecular findings among patients referred for clinical whole-exome sequencing. JAMA 312, 1870–1879 (2014)

c34. Trifunovic A, Wredenberg A, Falkenberg M, Spelbrink JN, Rovio AT, Bruder CE, Bohlooly-Y M, GidlÖf S, Oldfors A, Wibom R, Törnell J, Jacobs HT, Larsson N-G. Premature ageing in mice expressing defective mitochondrial DNA polymerase. Nature 429, 417–423 (2004)

c35. Ross JM, Coppotelli G, Hoffer BJ, Olson L. Maternally transmitted mitochondrial DNA mutations can reduce lifespan. Sci. Rep. 4: 6569; DOI:10.1038/srep06569 (2014).

c36. Niemann J, Johne C, SchrÖder S, Koch F, Ibrahim SM, Schultz J, Tiedge M, Baltrusch S. An mtDNA mutation accelerates liver aging by interfering with the ROS response and mitochondrial life cycle. Free Radical Biology and Medicine 102, 174–187 (2017).

c37. Roolf C, Kretzschmar C, Timmer K, Sekora A, Knöbel G, Murua Escobar H, Fuellen G, Ibrahim SM, Tiedge M, Baltrusch S, Müller S, Köhling R, Junghanss C. Polymorphism in Murine mtATP8 Gene Correlates with Decreased Reactive Oxygen Species in Aging Hematopoietic Cells. In Vivo. 30:751–760 (2016)

c38. Kretzschmar C, Roolf C, Timmer K, Sekora A, Knübel G, Murua Escobar H, Jaster R, Müller S, Fuellen G, Köling R, Junghanss C. Uncoupling protein 2 deficiency results in higher neutrophil counts and lower B-cell counts during aging in mice. Exp Hematol 44:1085–1091 (2016)

c39. Kretzschmar C, Roolf C, Timmer K, Sekora A, Knübel G, Murua Escobar H, Fuellen G, Ibrahim SM, Tiedge M, Baltrusch S, Jaster R, Köhling R, Junghanss C. Polymorphisms of the murine mitochondrial ND4, CYTB and COX3 genes impact hematopoiesis during aging. Oncotarget 7: 74460–74472 (2016)

c40. Hirose M, Schilf P, Lange F, Mayer J, Reichart G, Maity P, Jöhren O, Schwaninger M, Scharffetter-Kochanek K, Sina C, Sadik CD, Köhling R, Miroux B, Ibrahim SM. Uncoupling protein 2 protects mice from aging. Mitochondrion 30: 42–40 (2016)

c41. Schmeing TM, Voorhees RM, Kelley AC, Ramakrishnan V. How mutations in tRNA distant from the anticodon affect the fidelity of decoding. Nat. Struct. Mol. Biol. 18, 432–436 (2011).

c42. Trifunovic A, Larsson NG. Mitochondrial dysfunction as a cause of ageing. J Intern Med 263, 167–178 (2008).

c43. Finsterer J. Cognitive dysfunctions in mitochondrial disorders. Acta Neurol Scand 26, 1–11 (2012).

c44. Pekny M, Pekna M. Astrocyte reactivity and reactive astrogliosis: costs and benefits. Physiol Rev 94, 1077–1098 (2014).

c45. Allard S, Scardochio T, Cuello AC, Ribeiro-da-Silva A. Correlation of cognitive performance and morphological changes in neocortical pyramidal neurons in aging. Neurobiol Aging 33: 1466–1480 (2012)

c46. Posimo JM, Titler AM, Choi HJ, Unnithan AS, Leak RK. Neocortex and allocortex respond differentially to cellular stress in vitro and aging in vivo. PLoS One 8(3):e58596 (2013).

c47. Bach D, Pich S, Soriano FX, Vega N, Baumgartner B, Oriola J, Daugaard JR, Lloberas J, Camps M, Zierath JR, Rabasa-Lhoret R, Wallberg-Henriksson H, Laville M, Palacin M, Vidal H, Rivera F, Brand M, Zorzano A. Mitofusin-2 determines mitochondrial network architecture and mitochondrial metabolism. A novel regulatory mechanism altered in obesity. J Biol Chem 278, 17190–17197 (2003)

c48. Torraco A, Peralta S, Iommarini L, Diaz F. Mitochondrial diseases part I: Mouse models of OXPHOS deficiencies caused by defects on respiratory complex subunits or assembly factors. Mitochondrion 0, 76–91 (2015).

c49. Moreno-Loshuertos R, Ferrin G, Acln-Perez R, Gallardo ME, Viscomi C, Pérez-Martos A, Zeviani M, Fernández-Silva P, Enríquez JA. Evolution Meets Disease: Penetrance and Functional Epistasis of Mitochondrial tRNA Mutations. PLOS Genet 7, e1001379 (2011).

c50. O’Keefe J, Nadel L. The Hippocampus as a Cognitive Map. NY: Oxford University Press (1978).

c51. LØmo T. Frequency potentiation of excitatory synaptic activity in the dentate area of the hippocampal formation. Acta Physiologica Scandinavica 68 (Suppl 277), 128 (1966).

c52. Bliss TVP, Collingridge GL. A synaptic model of memory: long term potentiation in the hippocampus. Nature 361, 31–39 (1993).

c53. Kumar A. Long-Term Potentiation at CA3-CA1 hippocampal synapses with special emphasis on aging, disease and stress. Front Aging Neurosci. 3, 7 (2011).

c54. Knapp LT, Klann E. Role of reactive oxygen species in hippocampal long-term potentiation: contributory or inhibitory? J. Neurosci. Res. 70 (1), 1–7 (2002).

c55. Lynch MA. Long-term potentiation and memory. Physiol Rev. 84 (1), 87–136 (2004).

c56. Huang Y, Hu Z, Liu G, Zhou W, Zhang Y. Cytokines induced by long-term potentiation (LTP) recording: a potential explanation for the lack of correspondence between learning/memory performance and LTP. Neuroscience 231, 432–43 (2013).

c57. Lodato S, Arlotta P. Generating neuronal diversity in the mammalian cerebral cortex. Annu. Rev. Cell Dev. Biol. 31, 699–720 (2015).

c58. Serrano F, Klann E. Reactive oxygen species and synaptic plasticity in the aging hippocampus. Ageing Res Rev. 3 (4), 431–43 (2004).

c59. Ramos ES, Larsson N-G, Mourier A. Bioenergetic roles of mitochondrial fusion. Biochimica et Biophysica Acta 1857, 1277–1283 (2016).

c60. Chen H, McCaffery JM, Chan DC. Mitochondrial fusion protects against neurodegeneration in the cerebellum. Cell 130, 548–562 (2007).

c61. Thomas CT, Ogle SB, Rumney BM, May HG, Adelson PD, Lifshitz J. Does time heal all wounds? Experimental diffuse traumatic brain injury results in persisting histopathology in the thalamus. Behav. Brain Res., DOI:10.1016/j.bbr.2016.12.038 (2017).

c62. Daverey A, Agrawal SK. Curcumin alleviates oxidative stress and mitochondrial dysfunction in astrocytes. Neuroscience 333, 92–103 (2016).

c63. Zou X, Ratti BA, O'Brien JG, Lautenschlager SO, Gius D, Bonini MG, Zhu Y. Manganese superoxide dismutase (SOD2): is there a center in the universe of mitochondrial redox signaling? J Bioernerg. Biomembr., DOI 10.1007/s10863-017-9718-8

c64. Radford NB, Wan B, Richmann A, Szczepaniak LS, Li JL, Li K, Pfeiffer K, Schagger H, Garry DJ, Moreadith RW. Cardiac dysfunction in mice lacking cytochrome-c oxidase subunit VIaH. Am J Physiol Heart Circ Physiol. 282, H726–H733 (2002).

c65. Dell’agnello C, Leo S, Agostino A, Szabadkai G, Tiveron C, Zulian A, Prelle A, Roubertoux P, Rizzuto R, Zeviani M. Increased longevity and refractoriness to Ca(2+)- dependent neurodegeneration in Surf1 knockout mice. Hum Mol Genet. 16, 431–444 (2007) .

c66. Yang H, Brosel S, Acin-Perez R, Slavkovich V, Nishino I, Khan R, Goldberg IJ, Graziano J, Manfredi G, Schon EA. Analysis of mouse models of cytochrome c oxidase deficiency owing to mutations in SCO2. Hum Mol Genet. 19, 170–180 (2010).

c67. Diaz F, Thomas CK, Garcia S, Hernandez D, Moraes CT. Mice lacking COX10 in skeletal muscle recapitulate the phenotype of progressive mitochondrial myopathies associated with cytochrome c oxidase deficiency. Hum Mol Genet. 14, 2737–2748 (2005).

c68. Diaz F, Garcia S, Padgett KR, Moraes CT. A defect in the mitochondrial complex III, but not complex IV, triggers early ROS-dependent damage in defined brain regions. Hum Mol Genet. 21, 5066–5077 (2012).

c69. Yu X, Gimsa U, Wester-Rosenlöf L, Kanitz E, Otten W, Kunz M, Ibrahim S. Dissecting the effects of mtDNA variations on complex traits using mouse conplastic strains. Genome Res. 19, 159–165 (2009).

c70. Sachadyn P, Zhang XM, Clark LD, Naviaux RK, Heber-Katz E. Naturally occurring mitochondrial DNA heteroplasmy in the MRL mouse. Mitochondrion 8, 358–366 (2008).

c71. Chomyn A, Martinuzzi A, Yoneda M, Daga A, Hurko O, Johns D, Lai ST, Nonaka I, Angelini C, Attardi G. MELAS mutation in mtDNA binding site for transcription factor causes defects in protein synthesis and in respiration but no change in levels of upstream and downstream mature transcripts. Proc. Natl. Acad. Sci. 89, 4221–4225 (1992).

c72. Mayer J, Reichart G, Tokay T, Lange F, Baltrusch S, Junghanss C, Wolkenhauer O, Jaster R, Kunz M, Tiedge M, Ibrahim S, Fuellen G, Köhling R. Reduced spatial learning ability as consequence of elevated superoxide levels in complex I mouse mutants. PLoS One. 10(4); DOI:10.1371/journal.pone.0123863 (2015).

c73. Gil-Pagés M, Stiles RJ, Parks CA, Neier SC, Radulovic M, Oliveros A, Ferrer A, Reed BK, Wilton KM, Schrum AG. Slow angled-descent forepaw grasping (SLAG): an innate behavioral task for identification of individual experimental mice possessing functional vision. Behavioral and Brain Functions 9, 35 (2013).

